# Chaos Game Representation: An Alignment-Free Technique for Exploring Evolutionary Relationships of Protein Sequences

**DOI:** 10.1101/276915

**Authors:** Priyasma Bhoumik, Austin L. Hughes

**Affiliations:** Department of Biological Sciences, University of South Carolina, Columbia SC 29208

**Keywords:** Chaos game representation, viral evolution, phylogenetic trees, cluster analysis, alignment-free method

## Abstract

Chaos Game Representation (CGR) is an iterative mapping technique, which shows patterns in amino acids or nucleotide sequences. Here we present a method for using CGR to explore evolutionary relationships of protein sequences based on amino acid properties and illustrate the approach with complete sets of protein translations from viral genomes. In an analysis of complete polyprotein sequences from the viral family Flaviviridae, the CGR method was able to cluster members of major viral groups together, but relations within groups were not well resolved in comparison to an alignment-based phylogeny. We applied the method to members of five different families of ssRNA positive-strand viruses, and each family formed a distinct cluster. We also present a method of testing the reliability of clustering in CGR-based trees, which involves the use of multiple random starting points for the CGR process.

## Introduction

Chaos theory describes behavior of dynamical systems; that is, systems involving processes in motion ^3, 10^. Such systems evolve with time and are highly sensitive to initial conditions. Chaos theory involves study of the unstable and irregular complex patterns in order to find order and regularity from them ^3, 10^ since the formation of a complex pattern often has a very simple process responsible for it. These complex patterns are called fractals.^16, 18, 25^ Chaos game representation (CGR) is an iterative mapping technique ^1, 18^ derived from chaos theory, ^10^ which can be used to search for patterns in DNA sequences or amino acid sequences.^4, 5, 17, 18^ CGR is different from most methods of sequence analysis in that it provides the user with a visual image depicting both composition and sequentiality.^5, 11^ This technique has been utilized previously to obtain information about DNA and amino acid sequences of a variety of organisms.^8, 9, 18, 19^

Many widely used techniques for reconstructing phylogenetic relationships from molecular sequence data depend on sequence alignment.^15, 21, 23, 24^ However, alignment becomes difficult or unreliable in the case of distantly related sequences,^31–33^ even though evolutionary information may still be present. For these reasons, alternative methods of phylogenetic sequence reconstruction based on gene family content ^17^ or on gene order^22^ have been used in cases where complete genome sequences are available. Here we examine the potential of methods based on CGR for clustering amino acid sequences without alignment.^2^ Because of their high mutation rates, protein sequences of viruses often cannot be aligned except in the case of closely related virus species. Moreover, because most viruses have small genomes, methods based on gene family content or gene order may not be very informative. Thus, CGR may be useful in elucidating evolutionary relationships of more distantly related virus taxa and we illustrate application of the method using amino acid sequence data from viruses.

## Methods

### Chaos game representation

CGR or chaos game representation is an algorithm that uses iterations in order to generate a pattern by utilizing the nucleotides in DNA or amino acids in protein sequences. CGR assigns a coordinate value to each unit in a sequence and hence a characteristic visual pattern is generated for each sequence.^18^ In the case of a DNA sequences, CGR assigns each of the four possible nucleotides A, T, G and C to one of the four vertices of a square. In our study, we used protein sequences; the 20 amino acids were divided into 4 groups, and each of these groups (designated A, B, C and D) was assigned to one of the four vertices of the square. We used two methods of grouping: (1) random groups of five amino acids each (2) groups based on amino acid residue chemical properties (charge and polarity): A = D, E (negatively charged); B = K, R, H (positively charged); C = S, T, N, C, Y, Q (neutral/polar); D = G, A, V, L, I, M, P (neutral/non-polar)

In Cartesian space, the CGR vertices were assigned to the four groups of amino acids as follows: A = (0, 0); B = (0, 1); C = (1, 1); D = (1, 0). Successive points in the CGR were generated by an iterated function system defined by the following equations

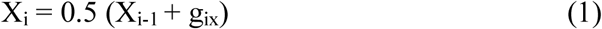

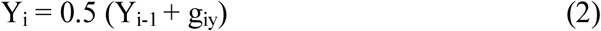

where g_ix_ and g_iy_ are the X and Y co-ordinates respectively, of the vertices corresponding to the amino acid residue at position *i* in the sequence. The coordinates for each amino acid residue were calculated iteratively using (0.5, 0.5) as an arbitrary starting position, and the pointer was moved half the distance to the next nucleotide to determine the next position.^1, 19, 14^ The output file contained x, y coordinate values for each amino acid present in the input sequence. These x and y coordinate values were plotted as scatterplots (Figure 1). To provide a test of the robustness of clustering patterns, 100 random starting points were chosen.

**Figure 1.**
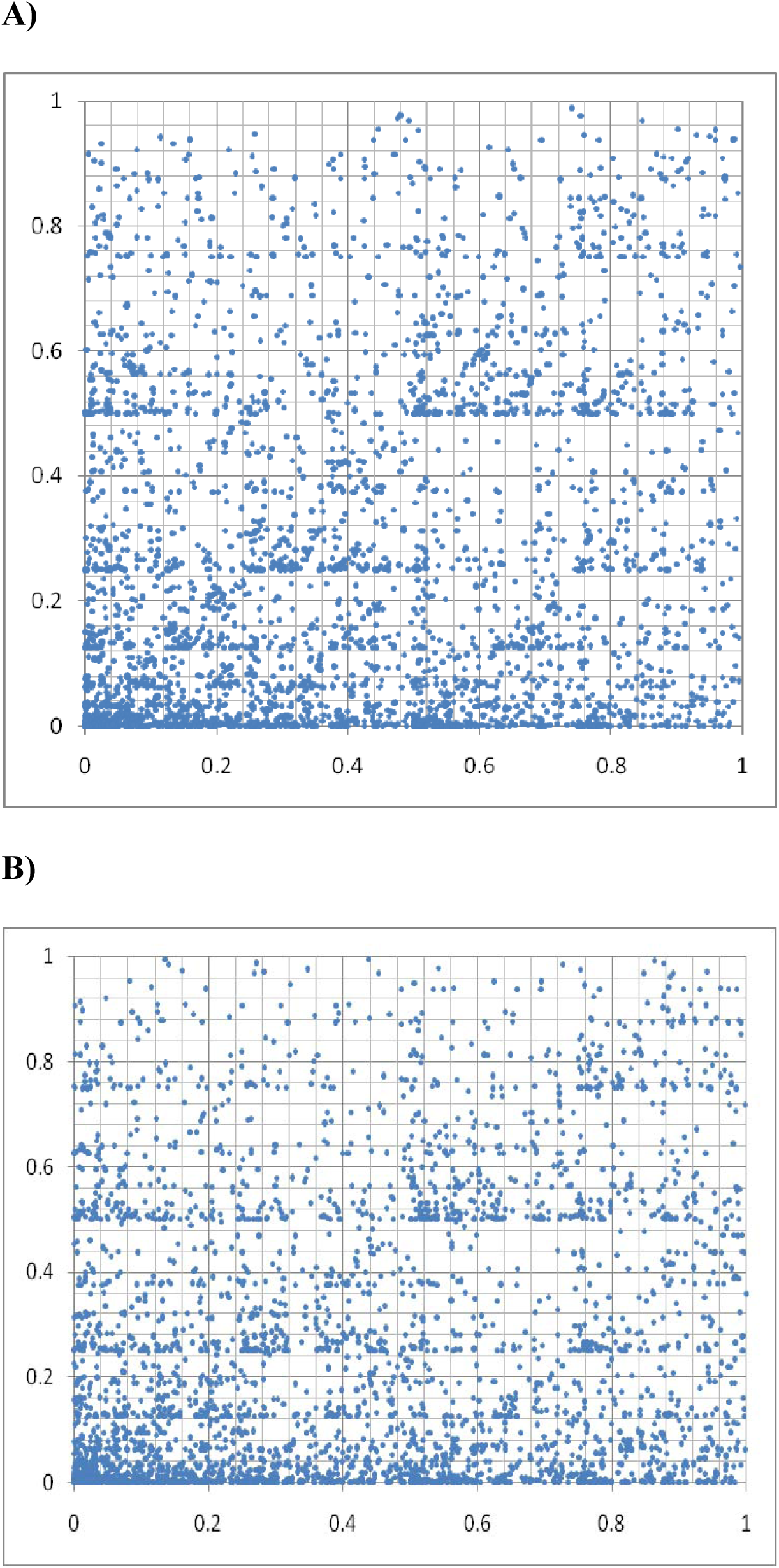
Examples of chaos game plots based on amino acid chemical properties: (A) Hepatitis C Virus 4 (HCV4) (B) Dengue Virus 1 (DENV1)

As in a previous study,^28^ each graph was divided into 4 quadrants; and the average x and y values of each point were calculated for each quadrant. A quadrilateral was then obtained from the 4 points corresponding to the average values for each quadrant; and the intersection point of the diagonals of the quadrilateral was identified. The evolutionary distance between two genomes was then defined as the Euclidean distance between the intersections of the diagonals of the quadrilaterals calculated for each of the two genomes. From the matrix of pairwise distances, a phylogenetic tree was constructed by the UPGMA method in Mega 4.^27^ In order to provide a test of the reliability of CGR trees based on amino acid residue chemical properties, trees were generated from random starting points in 100 trials.^13^ In the case of random grouping of the amino acids, random groups were created 100 times; the phylogenetic tree was constructed for each grouping. The consensus of the resulting 100 trees was constructed using Phylip Consense.^13^

Amino acid sequences of complete sets of proteins were obtained from the Genbank database. The accession numbers of all the sequences used in this study are listed (Table S1).

1. Flavivirus consisting of West Nile 1-2 (WNV), Yellow Fever (YFV), St.Louis Encephalitis (SLEV), Japanese Encephalitis (JEV) and Murray Valley Encephalitis (MVEV), Dengue 1-4 (DENV) considered as Flavivirus group of viruses (Flaviviridae family)
2. Hepacivirus^7^ consisting of HCV 1-6 considered as Hepacivirus group (Flaviviridae family) and Hepatitis E virus (HEPV - Hepeviridae family)
3. Human Rhinovirus (RV) B14, B27, Entero virus (EV) B1, C11 (Picornaviridae family)
4. Chikungunya virus (CHIKV), Ross River virus (RRV), Mayaro virus (MAYV) (Togaviridae family)
5. Bean Common Mosaic virus (BCMV), Bean Yellow Mosaic virus (BYMV), Potato virus Y (PVY), Potato virus V (PVV) (Potyviridae family)
6. Tomato Mosaic virus (ToMV), Tobacco Mild Green Mosaic virus (TMGMV), Tobacco Mosaic virus (TMV)

In the case of the Flaviviridae, we aligned complete polyprotein sequences by CLUSLAL X^29^ and constructed a tree based on the neighbor-joining (NJ) method^26^ with the Poisson-corrected amino acid distance using MEGA 4^27^. The reliability of clustering patterns in the tree was tested by bootstrapping;^12^ 100 bootstrap replicates were used.

## Examples

Figure 1 show examples of CGR plots using the groups based on amino acid residue chemical properties for complete polyprotein amino acid sequences of two flaviviruses, HCV4 and DENV4; the figure illustrates the distinct patterns produced by the two sequences. Figure 2 compares the consensus CGR tree of complete flavivirus polyproteins based on amino acid residue chemical properties (Figure 2A) with the NJ tree based on aligned sequences (Figure 2B). Two major patterns were similar between the two trees; namely, the HCV sequences clustered apart from the remainder of the flaviviruses, and the dengue viruses formed a distinct cluster (Figure 2). In the CGR tree, the cluster of HCV received 81% from 100 trials using different random starting points, while that of dengue viruses received 82% support (Figure 2A). However, the trees differed in a number of details. Most notably WNV1 and WNV2 grouped together in the sequence-based tree (Figure 2A), but in the CGR tree WNV1 clustered with YFV (Figure 2B). However, support for the latter patterns from random starting points was low (47%; Figure 2A). In general, bootstrap support values in the alignment-based tree were much higher than support values in the CGR tree based on trials using different random starting points (Figure 2). In fact, all bootstrap values in the alignment-based tree were 99% or greater except for those within the HCV cluster (Figure 2B).

**Figure 2.**
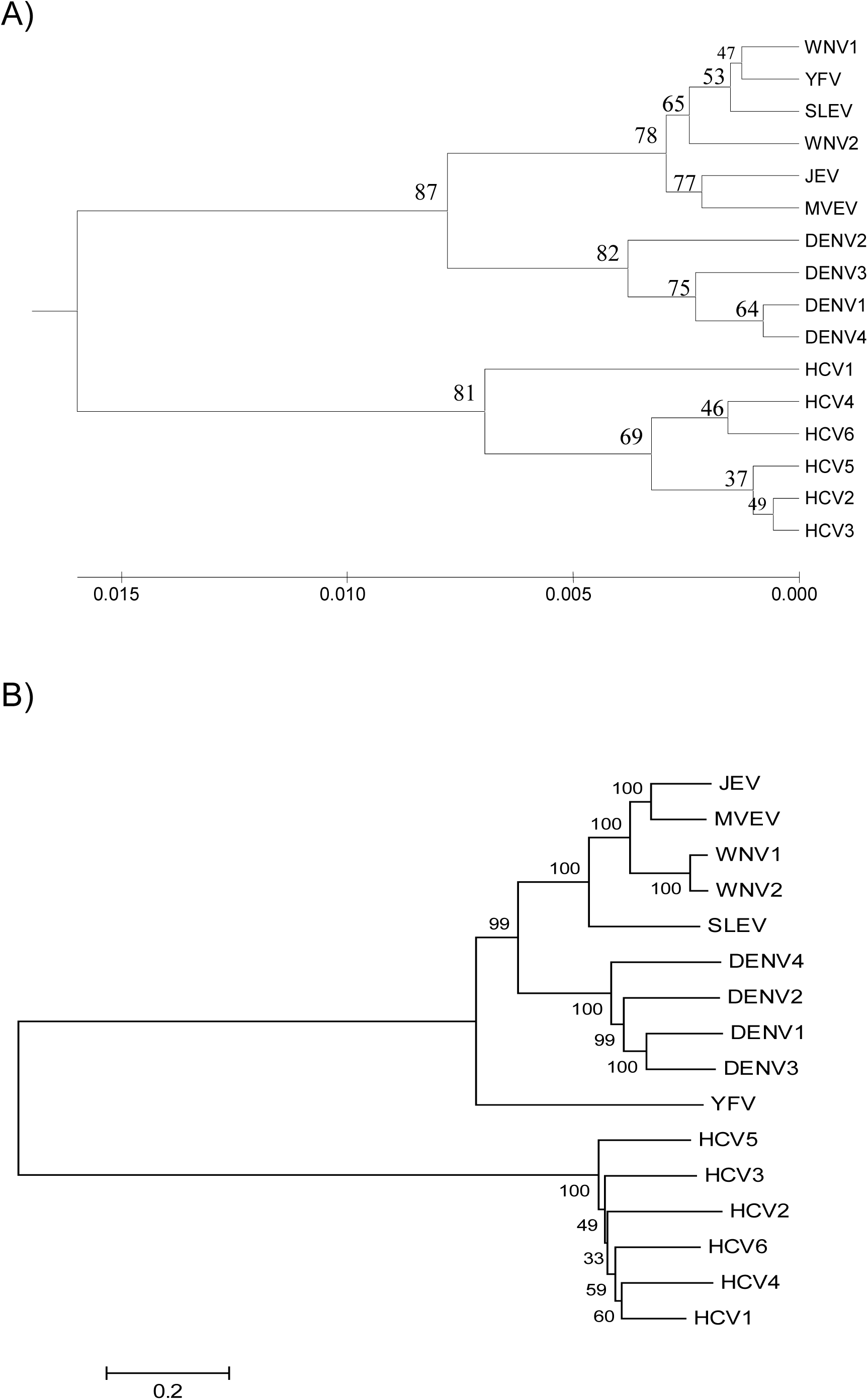
(A) CGR tree of flaviviruses based on amino acid chemical properties; numbers on the branches represent the percentage of 100 trials using different random starting points supporting the branch. (B) NJ tree based on aligned polyprotein sequences from the same viruses; numbers on the branches represent the percentage of 100 bootstrap samples supporting the branch.

In contrast to the CGR tree based on acid residue chemical properties, CGR trees based on random groupings of the amino acids provided little resolution of flavivirus polyprotein relationships (Figure 3). The HCV sequences grouped together (with 82% support from 100 different random groupings), but other biologically meaningful clusters were not resolved.

**Figure 3.**
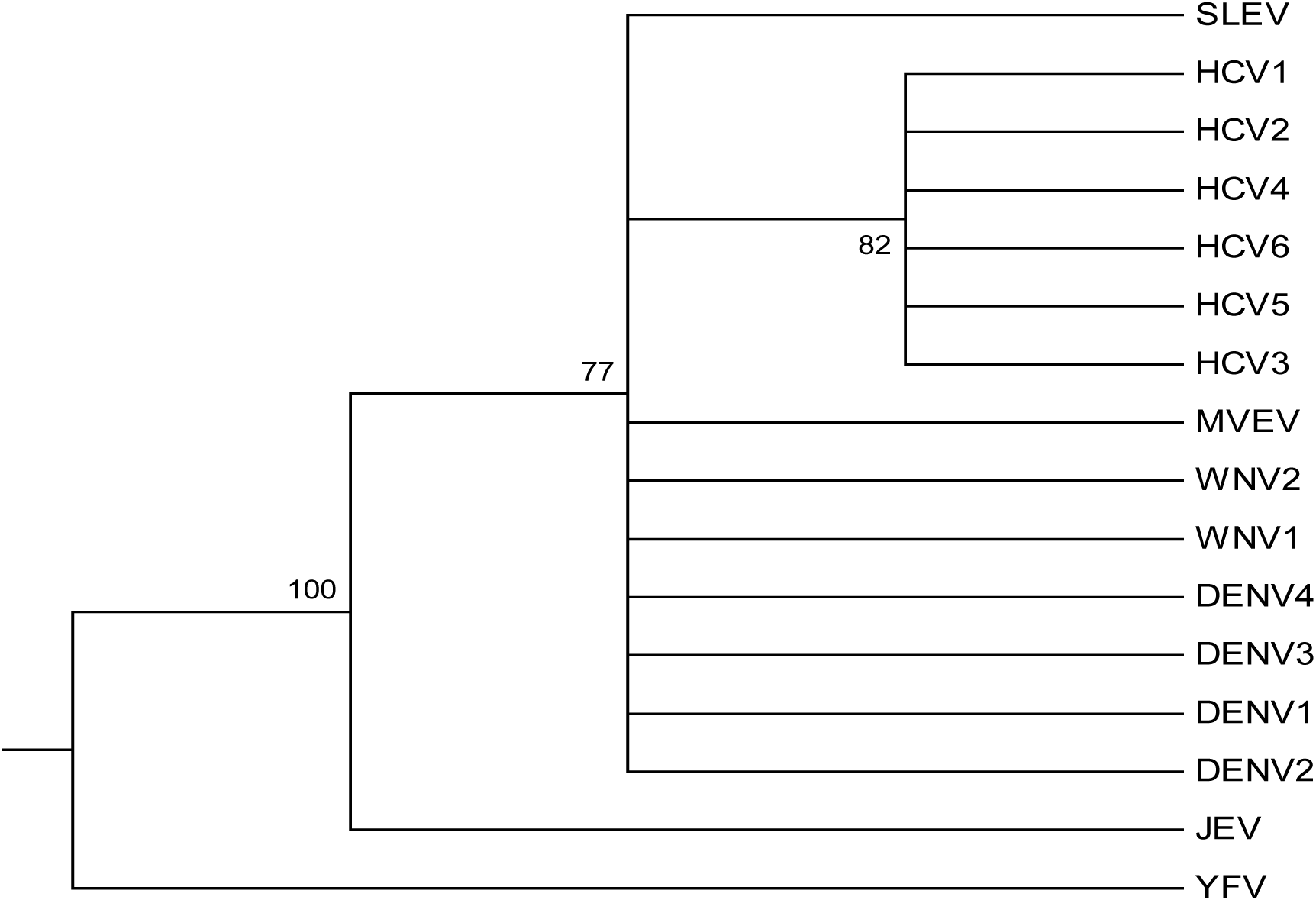
Consensus of 100 CGR trees of flaviviruses based on random amino acid groups.

In order to explore the utility of CGR for determining the relationships of widely divergent groups of viruses, we applied CGR analysis based on amino acid residue chemical properties to complete sets of protein sequences encoded by the genomes of 16 viruses representing the families Flaviviridae, Picornaviridae, Potyviridae, Togaviridae and Virgaviridae, all of which are ssRNA positive-strand viruses with no DNA stage.^30^ Each family formed a distinct cluster; and the support for family specific-clusters was 75% or better in every case except Virgaviridae (Figure 4). The Potyviridae and Virgaviridae, which are both families of plant viruses,^30^ clustered together with 77% support (Figure 4). The Picornaviridae, a family of animal viruses,^30^ formed a cluster with the latter two families that received 81% support.

**Figure 4.**
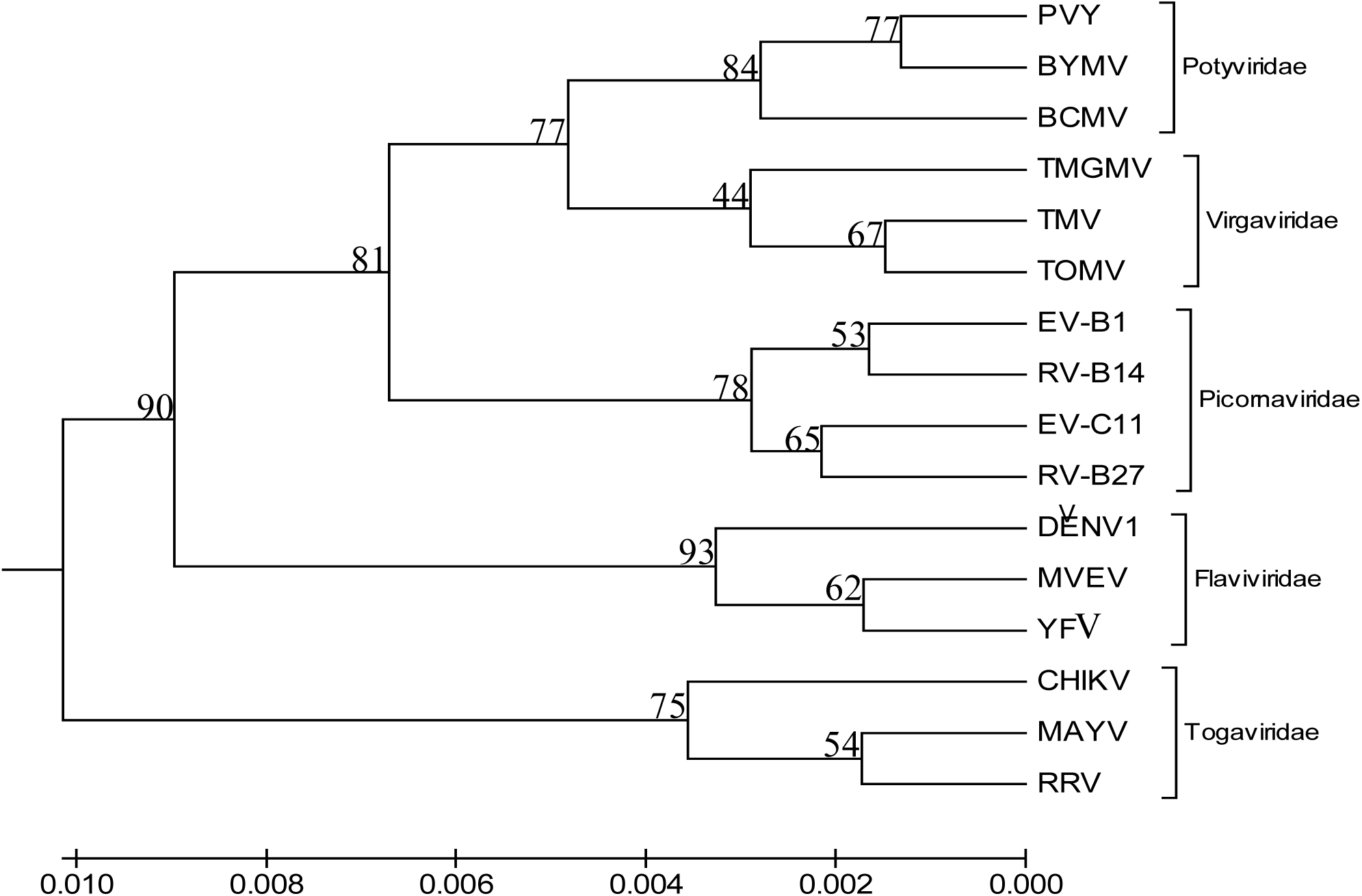
CGR tree of members of five families of ssRNA positive-strand viruses; numbers on the branches represent the percentage of 100 trials using different random starting points supporting the branch.

## Discussion

Application of CGR based on amino acid sequences to two sample datasets illustrated the potential strengths of this method for examining evolutionary relationship, as well as a number of weaknesses. First, we found that CGR analysis based on random amino acid groups provided little insight into evolutionary relationships (Figure 3). On the other hand, when the amino acids were grouped on the basis of chemical properties, the method performed much better. In the case of flavivirus polyproteins, we were able to compare the results of CGR analysis with a tree based on a sequence alignment. This comparison showed that the CGR method based on amino acid residue chemical properties captured broad patterns of relationship fairly well (Figure 2). However, more detailed relationships were apparently not very well captured by CGR, since there were numerous differences of detail between the CGR- and sequence-based trees (Figure 2).

A second example involved viruses belonging to five different families of ssRNA positive-strand viruses. This example provided further support for the ability of CGR to cluster related sequences, since each family formed a distinct cluster in the resulting tree (Figure 4). In addition, the CGR analyses of these families provided a hypothesis regarding their relationships, which are so far poorly understood. Attempts to resolve the higher relationships of these viruses have generally relied on the alignment of RNA-dependent RNA polymerase (RDRP) sequences.^6, 20, 33^ However, the alignment of RDRP amino acid sequences across different ssRNA virus families have proved challenging, and phylogenetic analyses have provided poor resolution of relationships.^6, 20, 33^ Our results were intriguing in that well-supported and potentially biologically meaningful clustering patterns were produced; for example, the clustering of the plant virus families Potyviridae and Virgaviridae. This pattern has not generally been seen in RDRP-based phylogenies, although these taxa clustered close to each other in the analysis of Bruenn.^6^

In this paper, we introduce a technique for testing the reliability of clustering patterns in CGR-based trees, involving repetitive trials of the CGR process multiple times using randomly chosen starting points. The percentages of trials supporting a given clustering pattern provides an index of the reliability of that pattern by analogy with the “bootstrapping” method commonly used to test clustering patterns in alignment-based trees.^12^ However, in the analysis of flaviviruses, the support values provided by trials using random starting points for CGR tended to be lower than the bootstrap values in the alignment-based tree (Figure 2). Thus, the former method appears to be more conservative than bootstrapping, and our results suggest that a reasonable cut-off for strong support in a CGR tree would be about 70% of trials.

In the present use of CGR, the algorithm used to generate the distance matrix captured only a small amount of the difference between any two CGR plots. The method might be improved in the future by using a more complex distance that incorporates all of the information in the CGR plot; for example, a multivariate approach that takes into account the distance between every point in one plot and its nearest neighbor in the other plot. Exploring more complex distances may provide a way of improving the method so that it captures relationships among closely related sequences as well as the broad patterns of relationship captured by the present method.

## Availability

The programs used and the user manual is available upon request via email.

## Supplementary Data

Supplementary Table S1: Accession numbers of viral sequences used in analyses.

